# PDLIM2 in Lung Adenocarcinoma Metastasis

**DOI:** 10.64898/2025.12.03.692242

**Authors:** Feng Gao, Yadong Xiao, Hongqiao Zhang, Steven D. Shapiro, Zhaoxia Qu, Gutian Xiao

## Abstract

Human and mouse studies have established the unique PDZ-LIM domain-containing protein PDLIM2 as a common tumor suppressor that is especially vital for suppressing the tumorigenesis and therapeutic resistance of lung cancer, the leading cause of cancer-related deaths among both men and women. However, the role of PDLIM2 in tumor metastasis, the predominant cause of cancer morbidity and mortality, is yet to be determined. Here, we report that PDLIM2 repression was positively associated with the metastasis of human lung adenocarcinoma, the major type of non-small cell lung cancer that accounts for more than 40% of all cases of human lung cancer. Interestingly, PDLIM2 repression was also correlated with oncogenic KRAS and/or TP53 mutations, two common drivers of human lung adenocarcinoma that often co-occur. In mice, in comparison to concurrently inducing mutant KRAS expression and TP53 deletion, additional co-ablation of PDLIM2 significantly increased the number and size of lung adenocarcinomas in the lung, and more importantly, the distant metastasis of lung tumor cells. The increased metastasis was accompanied by decreased anti-tumor immunity and increased pro-tumor inflammation. These data demonstrate the role of PDLIM2 in suppressing lung adenocarcinoma metastasis, thereby improving our understanding of this crucial tumor suppressor and lung cancer. They also provide a useful model for studying metastasis and testing new lung cancer treatments *in vivo*.

## Introduction

Lung cancer is the leading cause of cancer-related deaths among both men and women in the U.S. and worldwide [1]. Most lung cancer patients (up to 90%) die from metastasis, which is also the main reason for treatment failure. Lung cancer prognosis is very poor, with a single-digit 5-year survival rate for metastatic patients [1]. Hence, better understanding of lung cancer metastasis is important and urgent.

Mouse models provide a valuable tool for understanding tumor pathogenesis, testing new therapies, and identifying diagnostic and prognostic biomarkers [2]. The most used models of lung cancer are based on the conditional expression of the KRAS^G12D^ mutant or the administration of chemical carcinogens. KRAS mutations, including the KRAS^G12D^ mutation, are the most frequent oncogenic alterations in human lung cancer, and tobacco smoking, which contains urethane and many other chemical carcinogens, is the primary risk factor responsible for approximately 90% of all human lung cancers [3-9]. However, metastasis has not been found in those models, although they closely resemble the formation of human lung adenocarcinoma (LUAD), the most common type of non-small cell lung cancer (NSCLC) that makes up over 40% of all lung tumors. Co-deletion of TP53, a gene encoding the tumor suppressor p53 that is often mutated in human LUADs and particularly those harboring KRAS mutations, induces metastasis of murine LUADs driven by KRAS^G12D^ [9]. However, the distant metastasis rate in this model is low. To improve our understanding of lung cancer metastasis and combat this deadly disease, we need to develop new mouse models with better recapitulation of the high metastasis rate in patients.

In this regard, PDZ-LIM domain-containing protein 2 (PDLIM2, also known as Mystique or SLIM), a ubiquitously expressed protein with the highest level in the lung under normal conditions [10-12], is repressed in more than 90% of all NSCLC cases when 50% of the expression level of lung tissues adjacent to tumors is used as the cut-off [13, 14]. Moreover, PDLIM2 repression is associated with clinical outcomes negatively, such as poor patient survival and high resistance to the conventional chemotherapy and/or the innovative immune checkpoint inhibitors (ICIs) [14, 15]. Human cell line and mouse studies further demonstrate PDLIM2 repression as a causative driver of lung cancer and therapy resistance [13, 14]. For example, alveolar type 2 (AT2) epithelial cell-specific or global PDLIM2 deletion in mice is sufficient to promote primary LUAD that is highly chemo-resistant and completely resistant to ICIs [13, 14]. Notably, clinically feasible nano-delivery of exogenous PDLIM2 (nanoPDLIM2) or pharmacological induction of endogenous PDLIM2 shows a promising efficacy as a monotherapy, and in combination with chemo drugs and ICIs, completely eradicates all tumors in most animals without adding toxicity, in the urethane model of LUAD [13, 14, 16]. Of note, chemo drugs, ICIs, or their combinations fail to do so in a single animal in this autochthonous LUAD model. PDLIM2 deletion also significantly accelerates LUAD development in both KRAS^G12D^ and urethane models [13, 14]. Functionally, PDLIM2 has been shown to suppress tumor cell survival, proliferation, migration, invasion, and immune evasion [13, 14, 17].

However, the role of PDLIM2 in tumor metastasis has not been determined yet. This manuscript aims to address this scientifically and clinically important issue using mouse genetic models and next-generation sequencing data of human LUAD samples. Particularly, we will examine the relationships of PDLIM2 repression with LUAD metastasis and KRAS and/or TP53 mutations in humans and test if PDLIM2 co-deletion in KP mice increases LUAD distant metastasis. These studies will surely generate new knowledge on PDLIM2 and LUAD and develop a better *in vivo* model for LUAD basic and translational research, especially given the high rate of PDLIM2 repression in human LUAD, the sufficiency of PDLIM2 deletion alone in driving primary LUAD in mice, as well as the potential clinical applicability of nanoPDLIM2.

## Materials and Methods

### Human LUAD data

Large-scale single-cell RNA-sequencing (scRNA-seq) data of human LUAD were downloaded from NCBI BioProject #PRJNA591860 [18]. All codes used to generate the normalized cell expression data can be found on GitHub (czbiohub/scell_lung_adenocarcinoma and czbiohub/cerebra) and were described in the original paper. Cells of interest were extracted by tumor type (primary/metastatic), cell type (tumor/fibroblast/macrophage), and feature (PDLIM2). PDLIM2 expression levels in primary versus metastatic tumors at the tissue and cell levels were then analyzed, compared, and visualized with dot plots, where cells with the baseline of PDLIM2 expression were dismissed [19-21].

Human LUAD bulky RNA- and DNA-seq data from The Cancer Genome Atlas (TCGA) were analyzed to compare and visualize PDLIM2 expression levels in tumors with or without KRAS and/or TP53 mutations using cBioPortal [22].

### Animals and LUAD induction

PDLIM2^flox/flox^ mice under a pure FVB/NJ background have been described before [14]. KP (Lox-Stop-Lox-K-Ras^G12D^/p53^flox/flox^) mice were purchased from The Jackson Laboratory (Strain # 032435) and backcrossed more than ten generations to the pure FVB/N background. The two strains were then bred to generate PKP (PDLIM2^flox/flox^/Lox-Stop-Lox-K-Ras^G12D^/p53^flox/flox^) mice. Female KP and PKP mice of 6-8 weeks old were instilled intranasally 1 × 10^7^ plaque-forming units (pfu) of Cre-expressing adenovirus (AdenoCre) once per week for 6 weeks [3, 4, 14]. Eight- or sixteen-week post-AdenoCre instillation, all mice were euthanized for tumor examinations. Mouse tissues, including primary lung tumors and adjacent lung tissues, metastatic tumors, and lymph nodes (LNs), were excised and used for the assays below. All animals were maintained under a specific pathogen-free condition and used according to protocols approved by the University of Southern California (USC) IACUC.

### Enumeration and burden calculation of tumors in the lung

Images from both sides of each lung lobe were taken and imported into ImageJ software and deconvolved to define and trace the outline of tumors [3-8, 13, 14]. Tumors in the mouse lungs were counted. Since tumors are predominantly well-rounded spheres, the diameter of each tumor was therefore calculated, and the volumes of all tumors in each mouse were summed up and averaged.

### Histological and immunostaining assays

Tissues from mice were fixed in formalin, embedded in paraffin, and cut into 4-5μm thick sections. The tissue sections were deparaffinized, rehydrated, and stained with hematoxylin and eosin (H&E) or subjected to antigen retrieval (heated in citric acid buffer), Triton-100 permeabilization, serum blocking, sequential incubations with primary antibodies against surfactant protein C (SPC, a specific marker for lung AT2 epithelial cells) (Chemicon International) and fluorophore-conjugated secondary antibodies (Abcam), and nuclei counterstaining with 4′,6-diamidino-2-phenylindole (DAPI) [23-29]. Stained sections were mounted for imaging under a Leica DMi8 fluorescent microscope.

### Distant metastasis determination

In addition to anatomically distant tumors that were stained positively with SPC, distant metastasis in KP and PKP mice was also based on the positive SPC staining of accessory axillary (1), proper axillary (1), cervical (1), and inguinal (2) LNs.

### Flow cytometry analysis

Lung tumors and their adjacent lung tissues were minced, dispersed, and followed by digestion with a mix of collagenase and dispase on an orbital shaker [30]. Following the enzymatic and mechanical dissociation, the suspension was filtered, and red blood cells were lysed. The single-cell suspension was then subjected to Flow cytometry analysis [31-35]. Briefly, after blocking with anti-CD16/CD32, the cells were incubated with fluorescent-conjugated antibodies against the cell surface proteins on myeloid or lymphocyte cells. For lymphocyte cell analysis, following cell surface protein staining the cells were fixed with paraformaldehyde (2%), permeabilized with saponin (0.5%), and incubated with fluorescent conjugated antibodies against the intracellular protein Foxp3, a transcription factor that serves as a master regulator and a specific marker of CD4 regulatory T cells (Tregs). After a final wash, the antibody-stained cells were resuspended and run by BD Symphony A1 Flow cytometer (BD Biosciences). Data were acquired and analyzed by FlowJo software.

### Statistics

Student’s *t* test (2-tailed, unpaired) was used to assess the significance of differences between 2 groups. An ordinary 1-way ANOVA was used to assess the significance of differences among groups of more than 2 [36]. All bars in the figures represent mean ± SEM. P values less than 0.05 were considered statistically significant.

## Results

### Increased PDLIM2 repression in human metastatic LUAD

Previous analysis of the TCGA database has revealed that PDLIM2 is repressed in more than 90% of all human NSCLCs and in particular LUADs, and that the extent of PDLIM2 repression is associated with tumor progression [13, 14]. However, the TCGA database is mainly comprised of primary tumors. To compare PDLIM2 expression in primary and metastatic LUADs, we analyzed the NCBI BioProject #PRJNA591860 database, which contains scRNA-seq data of 11 primary and 21 metastatic LUADs. As expected, PDLIM2 expression was significantly lower in metastatic lung tumors in comparison to primary tumors (Fig.1A). Interestingly, PDLIM2 was further repressed not only in metastatic cells but also in their associated cells, particularly fibroblasts and macrophages, though the repression extents in those surrounding cells were lower than that in tumor cells (Fig.1B-E). These human data indicate that PDLIM2 repression in both tumor cells and tumor microenvironment is associated with primary LUAD progression to metastasis.

**Fig. 1.**
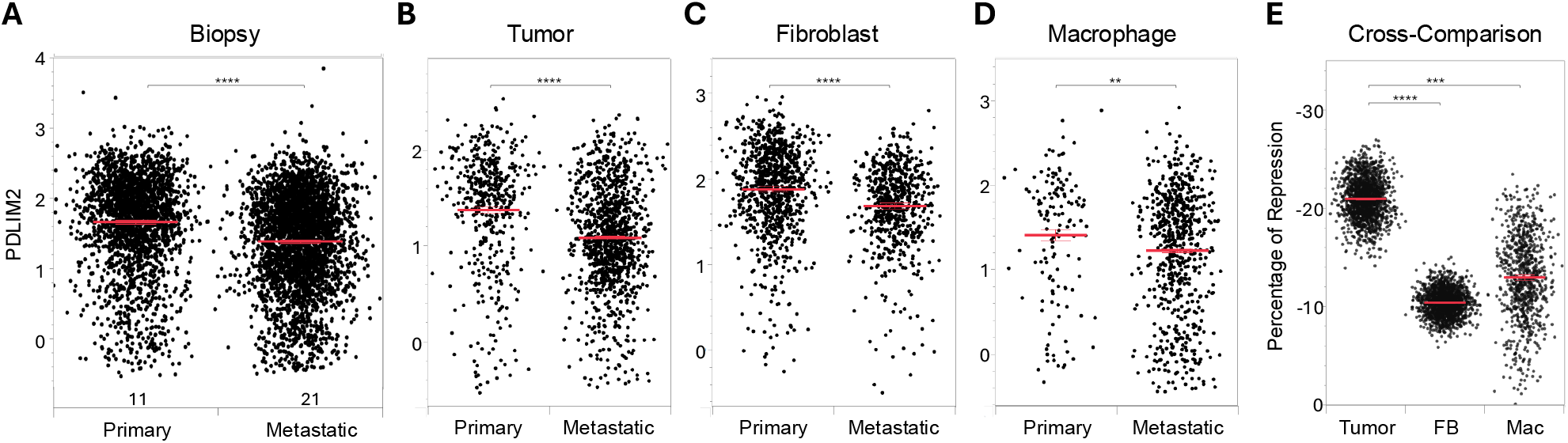
PDLIM2 is further repressed in human metastatic LUAD. **(A**. Analysis of NCBI BioProject #PRJNA591860 showing lower PDLIM2 expression in human metastatic LUAD. Patient sample numbers are listed under the dot plots. Each dot represents one cell. **B**. Lower PDLIM2 expression in human metastatic LUAD cells. **C**. Lower PDLIM2 expression in fibroblasts within human metastatic LUAD tissues. **D**. Lower PDLIM2 expression in macrophages within metastatic LUAD tissues. **E**. Cross comparison of PDLIM2 repression in tumor, fibroblast (FB), and macrophage (Mac) cells from bootstrapping calculation. The repression was calculated as the percentage change in PDLIM2 mean expression between the metastatic and primary cells within each group. Each dot represents one bootstrapping iteration. Data represent means ± SEM. ** P < 0.01; *** P < 0.001; **** P < 0.0001.

### Positive association of PDLIM2 repression with KRAS and TP53 mutations in human LUAD

Given the roles of KRAS and TP53 mutations in LUAD progression, we analyzed the TCGA database, because it contains both gene expression and mutation data. Of 500 human LUADs with known KRAS and TP53 mutation status in the database, 147 were wildtype for both genes, 100 harbored KRAS but not TP53 mutations, 200 had mutations in TP53 but not KRAS, and 53 carried mutations in both genes (Fig. 2). It is worth noting that the frequencies of KRAS and TP53 mutation alone and concurrently (20%, 40% and 10.6%, respectively) in LUADs were consistent with general predication [37]. Remarkably, compared to its level in tumors with wild-type KRAS and TP53, PDLIM2 was expressed significantly lower in KRAS and/or TP53 mutant tumors, with the lowest in those having KRAS and TP53 double mutants. These data suggest that PDLIM2 repression may play more significant roles in LUADs with mutations in KRAS and TP53, especially those with their co-occurring mutations.

**Fig 2.**
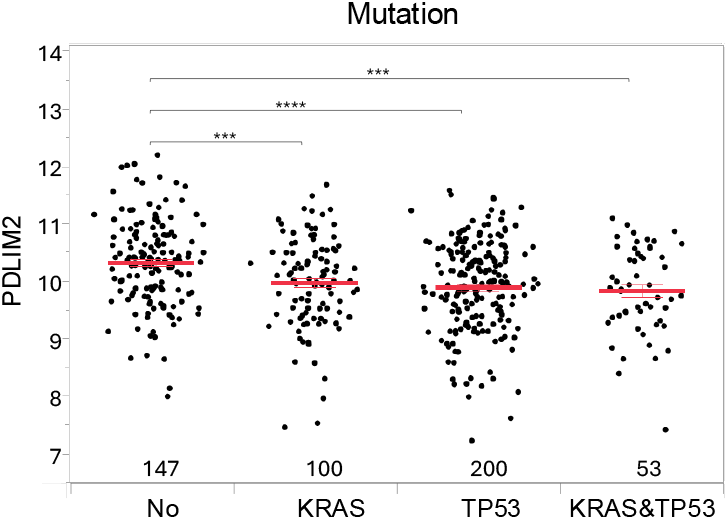
PDLIM2 repression is associated positively with KRAS and TP53 mutations in human LUAD. Analysis of the TCGA database showing a decreased PDLIM2 expression in human LUADs carrying KRAS and/or TP53 mutations versus those with wild-type KRAS and TP53. Each dot represents one patient, and patient sample numbers are listed under the dot plots. Data represent means ± SEM. *** P < 0.001; **** P < 0.0001.

### Increased LUAD formation and local lung metastasis in KP mice by PDLIM2 co-deletion

The human studies above prompted us to test whether PDLIM2 deficiency increases LUAD metastasis in the KP mouse model, in which KRAS mutant expression and TP53 genetic deletion can be induced simultaneously in the same lung cells by intranasal instillation of AdenoCre. To this end, we generated PKP mice in which PDLIM2 ablation, together with KRAS mutant expression and TP53 deletion, can be co-induced by the same method. As expected, all KP and PKP mice developed tumors in their lungs 8- or 16-weeks post-AdenoCre treatment (Fig.3A, and SFig.1). Histological studies indicated that those tumors were LUADs, as evidenced by their morphology and positive staining of SPC (Fig.3B). Note, SPC^+^ AT2 epithelial cells are the main cells-of-origin of LUAD [38-40].

**Fig 3.**
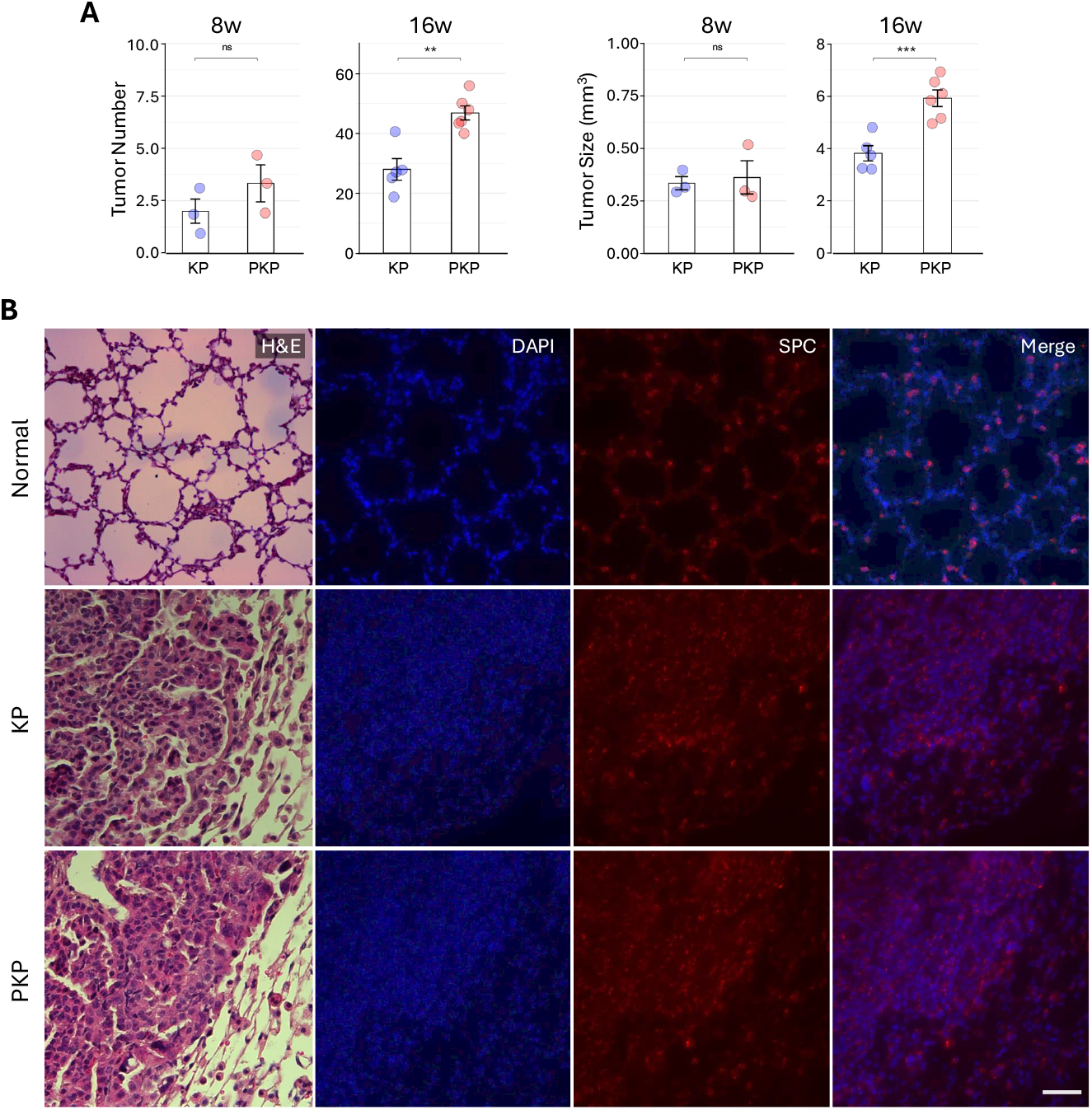
PDLIM2 co-deletion increases LUAD formation and local lung metastasis in KP mice. **A**. Increases tumor number and tumor size in the lung of PKP mice compared to KP mice 8 and 16 weeks post-AdenoCre administration. Data represent means ± SEM in A and B. ** P < 0.01; *** P < 0.001; ns: not significant. **B**. Histological studies showing LUAD morphology and positive SPC staining of tumors in the lungs of both KP and PKP mice. Scale bar: 50μm.

At the 8-week time point, no statistical difference in tumor number or size was found between KP and PKP mice. Moreover, both mice only had very few tiny tumor lesions in their lungs, implying no local lung metastasis in either of the mice at this early time point. At the 16-week time point, many more and much bigger tumors were seen in the lungs. However, tumor number and size were significantly higher in PKP mice compared to KP mice at this late time point. These data suggest that LUADs in PKP mice are more aggressive and with higher local lung metastasis.

### Increased LUAD distant metastasis in KP mice by PDLIM2 co-deletion

Notably, tumors were also found outside the lungs in PKP mice at the 16-week timepoint (Fig. 4A). The distant tumors were metastatic LUADs, because they were SPC-positive (Fig. 4B). Given the way-station role of LNs in distant metastasis, we examined whether there were LUAD cells within LNs at various locations, such as axilla, cervical, and inguinal. Remarkably, SPC-positive cells were detected in all PKP mice and often within multiple LNs of the same mouse (Fig. 4C, D). However, only 28.6% of KP mice had SPC-positive cells in their LNs. Moreover, fewer metastatic LUAD cells were found in fewer LNs in those KP mice. Taken together, these data demonstrate a crucial role of PDLIM2 in suppressing LUAD progression, local and distant metastasis.

**Fig 4.**
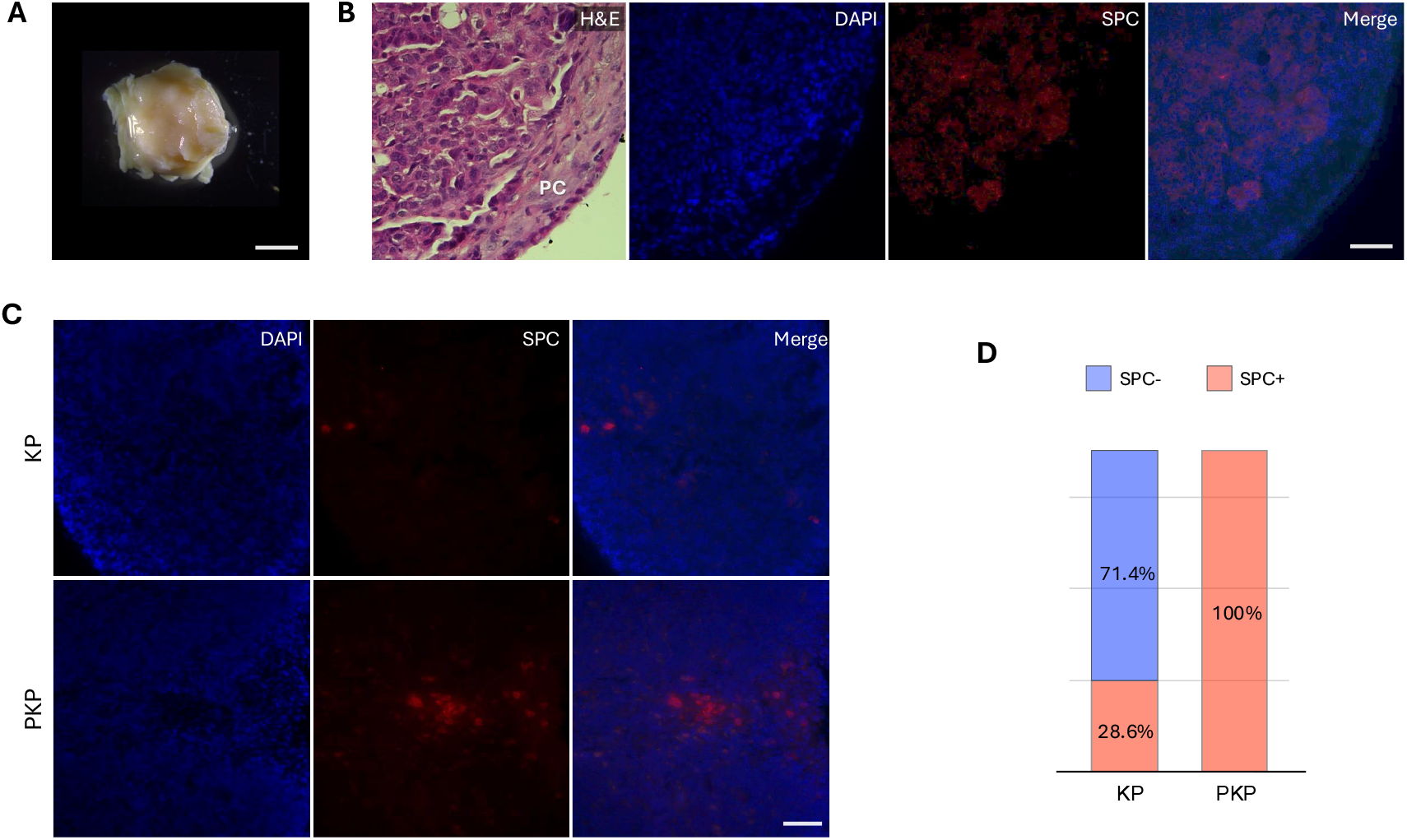
PDLIM2 co-deletion increases LUAD distant metastasis in KP mice. **A**. Representative metastatic tumor in PKP mice 16 weeks post-AdenoCre administration. Scale bar: 3mm. **B**. Histological studies showing the morphology and positive SPC staining of the metastatic LUAD tumors. **C**. Histological studies showing the positive SPC staining of the metastatic LUAD tumor cells in the LNs of KP and PKP mice 16 weeks post-AdenoCre administration. **D**. Distant metastasis rate in KP and PKP mice 16 weeks post-AdenoCre administration.

### Decreased antitumor immunity and increased pro-tumor inflammation by PDLIM2 co-deletion

To define how PDLIM2 co-deletion promotes LUAD progression and particularly from local to distant metastasis, we analyzed the immune profiles within tumors and surrounding tissues from KP and PKP mice. The interaction between tumor cells and immune cells plays a pivotal role in tumor progression and metastasis. Our flow cytometry analysis (SFig.2-4) revealed a higher ratio of total myeloid cells but a lower ratio of total lymphocytes in both tumors and surrounding tissues of PKP mice in comparison to KP mice, though being more evident in tumors (Fig. 5A-C). Neutrophils and particularly those expressing the immune checkpoint PD-L1 were significantly increased in the PKP mice (Fig. 5D-F). The main function of PD-L1 is to prevent T cells from killing tumor cells by binding to the PD-1 receptor on T cells. These data were highly consistent with that myeloid-to-lymphoid ratio, especially neutrophil-to-lymphocyte ratio (NLR), is a clinical biomarker for increased metastasis risk and poor prognosis. Macrophages and particularly alveolar macrophages (AMs), the most abundant immune cells in normal lungs and tumor microenvironments, were also increased in the PKP mice (Fig. 5G, H). In line with previous studies [33], all AMs of KP or PKP mice expressed PD-L1 on the surface (SFig.3). Remarkably, PD-L1’s levels on the AMs of PKP mice were significantly higher (Fig. 5I). Monocytic or granulocytic myeloid-derived suppressor cells (M- and G-MDSCs), both of which are tumor-promoting myeloid cells, were increased in the PKP mice as well (Fig. 5J, K). In contrast, monocytes were decreased in the PKP mice (Fig. 5L). It is plausible that the decrease in monocytes in the PKP mice was due to increased differentiation into macrophages and M-MDSCs from those precursor cells.

**Fig. 5.**
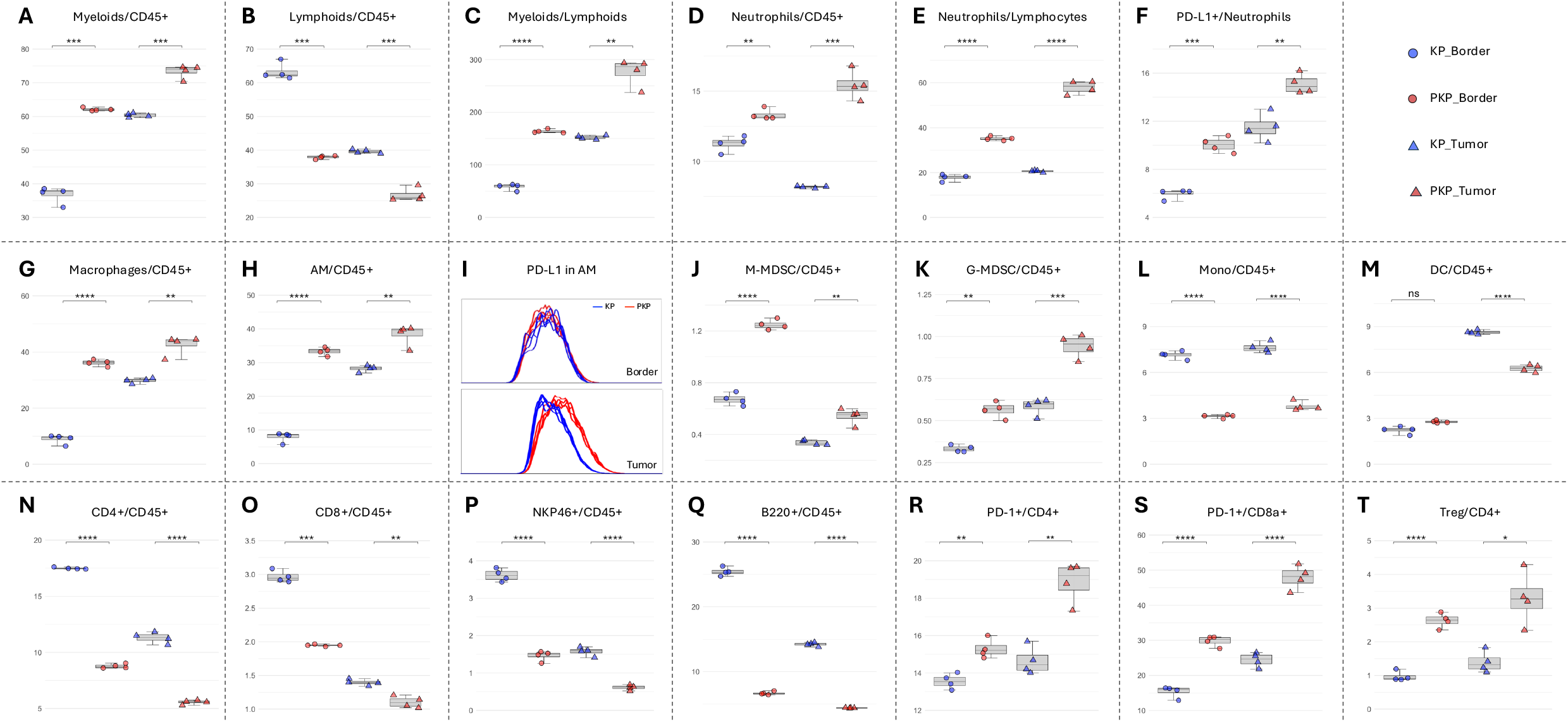
PDLIM2 co-deletion decreases antitumor immunity and increases pro-tumor inflammation. Flow cytometry analysis showing (**A**) increased myeloid cells, (**B**) decreased lymphocytes, (**C**) increased myeloid-to-lymphoid ratio, (**D**) increased neutrophils, (**E**) increased neutrophil-to-lymphocyte ratio, (**F**) increased PD-L1-postive neutrophils, (**G**) increased macrophages, (**H**) increased AMs, (**I**) increased PD-L1 on AMs, (**J**) increased M-MDSCs, (**K**) increased G-MDSCs, (**L**) decreased monocytes, (**M**) decreased DCs, (**N**) decreased CD4 T cells, (**O**) decreased CD8 T cells, decreased NK cells, (**Q**) decreased B cells, (**R**) increased PD-1-positive CD4 T cells, (**S**) increased PD-1-positive CD8 cells, and (**T**) increased CD4 Tregs in the LUADs and their surrounding lung tissues of PKP mice 16 weeks post-AdenoCre administration. Each line (I) or dot (others) represents one mouse. Data represent means ± SEM. * P < 0.05; ** P < 0.01; *** P < 0.001; **** P < 0.0001.

In line with their overall antitumor functions, the total numbers of dendritic cells (DCs), T cells, natural killer (NK) cells, and B cells were decreased in both tumors and the surrounding tissues of PKP mice in comparison to KP mice (Fig. 5M-Q). Furthermore, T cells in the PKP mice were more exhausted, as evidenced by significantly more CD4 and CD8 T cells expressing the immune checkpoint PD-1, a key exhaustion marker of T cells (Fig. 5R, S). Notably, the tumor-promoting CD4 Tregs were increased in the PKP mice (Fig. 5T). It is also noteworthy that compared to their surrounding ‘normal’ tissues, tumor tissues contained more tumor-promoting but fewer antitumor immune cells in both KP and PKP mice, supporting the common notion on the hijack and manipulation of immune cells by tumor cells for their own benefits and immune evasion. Hence, PDLIM2 repression further decreases antitumor immunity and increases pro-tumorigenic inflammation, enhancing LUAD progression and metastasis.

## Discussion

The studies above provide the first evidence demonstrating the crucial role of PDLIM2 in suppressing LUAD metastasis, besides its well-established tumor suppressor role in various cancers, and particularly in LUAD [13-15, 41-46]. As a matter of fact, they are the first investigation on the role of PDLIM2 in metastasis, at least under pathophysiological conditions. They also offer the first line of data linking PDLIM2 repression to KRAS and/or TP53 mutations and generate an ideal *in vivo* model for basic and translational research of LUAD.

LUADs in KP mice are the most famous lung cancer model with metastasis. However, the distant metastasis rate in this model is lower than 29% and needs to be improved significantly. Our PKP mice overcome the challenge with 100% penetrance. Of note, PDLIM2 is co-repressed in 100% of human LUADs with KRAS and TP53 mutations, indicating a strong human relevance for our model. It should also be pointed out that individual induction of KRAS mutation, PDLIM2 deletion, or TP53 ablation alone is sufficient to cause primary LUADs in mice, but all of them alone cannot induce LUAD metastasis [13, 14, 47].

PDLIM2 exerts its tumor-suppressor role via multiple mechanisms [17]. Perhaps, the most important is the negative regulation of NF-κB and STAT3, two pro-tumorigenic transcription factors that play central roles in inflammation, metabolic reprogramming, cell growth, tumor pathogenesis, and therapy resistance [14, 48-56]. PDLIM2 acts as a unique ubiquitin ligase enhancer (E5) to stabilize and chaperone the ubiquitin ligase (E3) SCF/β-TrCP to ubiquitinate nuclear RelA proteins for proteasomal degradation, thereby limiting NF-B activity [57, 58]. RelA, also known as p65, is the prototypical member of NF-κB that is persistently activated in a large plethora of cancers, including LUAD [14, 59-64]. Most likely, it uses the same or a similar mechanism to ubiquitinate and degrade nuclear STAT3. PDLIM2 repression in precancerous and cancer cells will therefore lead to uncontrolled proliferation and survival, increased migration and invasion, and immune evasion (through decreasing antigens and shifting immunity from antitumor into pro-tumor).

In addition to suppressing precancerous and cancer cells intrinsically, PDLIM2 also serves as a potent extrinsic tumor suppressor, particularly in macrophages, the most abundant immune cells in the tumor microenvironment. In human LUAD, PDLIM2 is repressed in tumor cells and associated macrophages (TAMs). Furthermore, the extent of PDLIM2’s repression in either tumor cells or TAMs is positively correlated with tumor progression, metastasis, and poor patient survival [14, 65]. In mice, deletion of PDLIM2 in AMs, like the nullification of AT2 epithelial-PDLIM2, promotes LUADs [14, 65]. Selective PDLIM2 loss in AMs and myeloid cells switches this central determinant of immunity from antitumor into pro-tumor phenotypes [65].

Intriguingly, PDLIM2 repression in lung cancer cells further reduces PDLIM2 in lung macrophages, and vice versa. This reciprocal repression involves PDLIM2’s transcriptional suppression by reactive oxygen species (ROS)-activated BACH1, a transcription repressor [65]. Tumor cells and macrophages are both major sources of ROS, which act in both autocrine and paracrine manners. Accordingly, ROS inhibitors can restore PDLIM2 expression and prevent lung cancer *in vivo* [65]. Given the relationship among tobacco smoking, NF-κB/STAT3 activation, ROS synthesis, and cancer pathogenesis [66-68], it is highly presumable that smoking activates NF-κB/STAT3 in lung epithelial cells and macrophages to induce ROS generation, leading to BACH1repression of PDLIM2 in those cells. PDLIM2 repression in turn results in persistent NF-κB/STAT3 activation and ROS production, transforming epithelial cells and hijacking immune cells. Chronic ROS exposure may also induce PDLIM2 epigenetic silencing and/or loss of heterozygosity (LOH) and cause alterations and/or genetic mutations in KRAS, TP53, and other genes in lung epithelial/tumor cells. Logically, KRAS and TP53 mutations may further contribute to PDLIM2 repression. These changes together drive lung cancer formation, progression, and metastasis.

In summary, our study establishes PDLIM2’s role in suppressing LUAD metastasis and its crosstalk with KRAS and TP53 mutations. They provide a rationale to target PDLIM2 to prevent and treat the metastasis of LUADs with mutated KRAS and TP53, particularly given the high therapeutic efficacy of the clinically feasible nanoPDLIM2 in the primary LUAD model. They also offer an *in vivo* model recapitulating the full process of human LUAD, from tumor initiation to progression to distant metastasis. They are thus important to the LUAD field and also to tumor metastasis at large, considering PDLIM2 repression as a common mechanism in human cancers.

## Author Contributions

F.G. performed and analyzed all assays and prepared all figures. Y.X. contributed to mouse breeding and colony maintenance and provided technical assistance. H.Z. assisted experimental studies. S.D.S provided constructive advice and feedback. Z.Q. and G.X. conceived and designed the study, led and contributed to all aspects of the analysis, and wrote the manuscript.

## Acknowledgments

We thank other team members for their suggestions and assistance.

## Ethics approval and consent to participate

All animal studies were approved by the Institutional Animal Care and Use Committee of the University of Southern California and carried out in accordance with NIH guidelines on animal care.

## Supplementary material

The supplementary material for this article can be found online.

## Supplementary figures legends

**SFig. 1.**
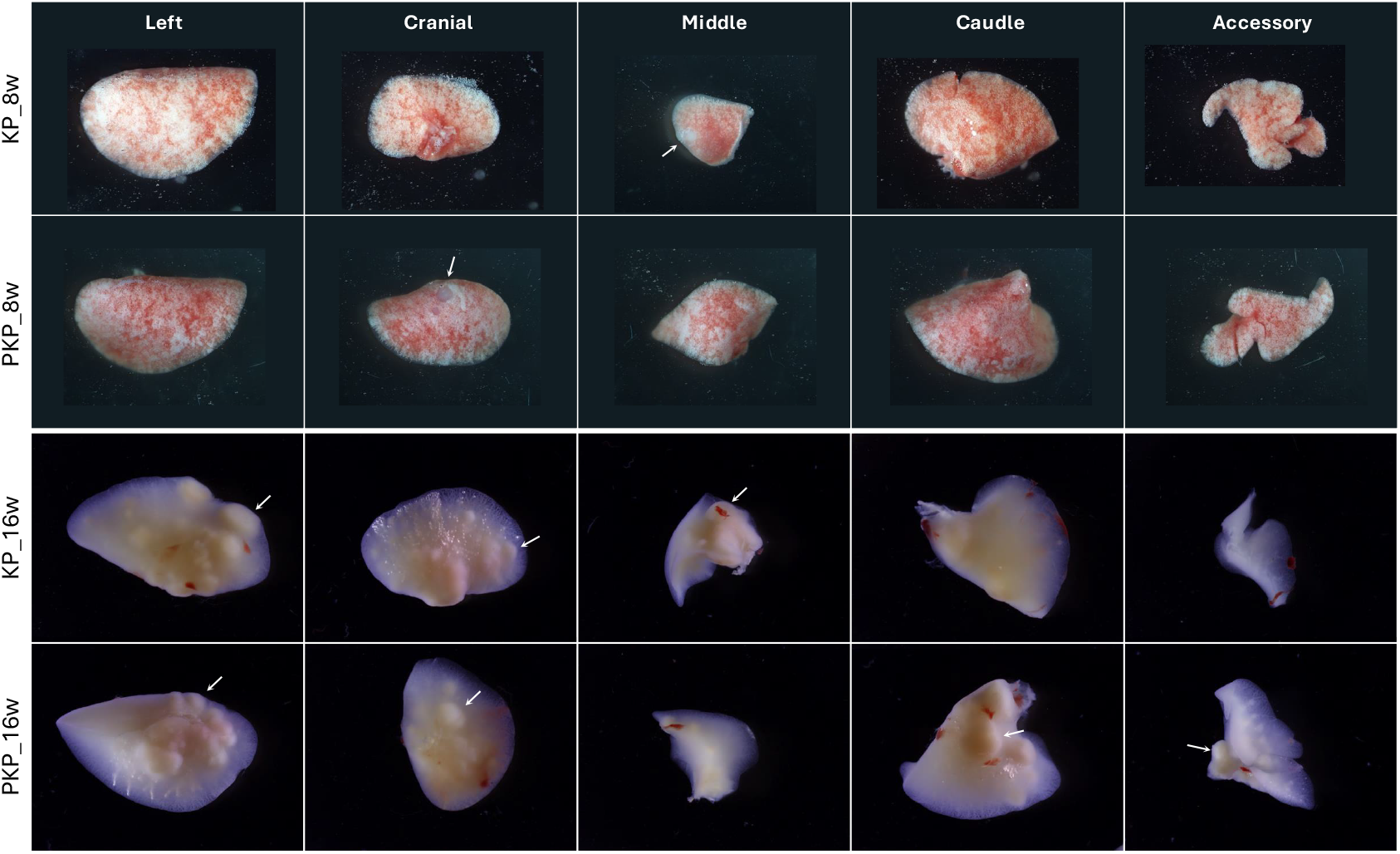
Representative lung tissue of KP and PKP mice 8- or 16-weeks post-AdenoCre administration. Scale bar: 3mm.

**SFig. 2.**
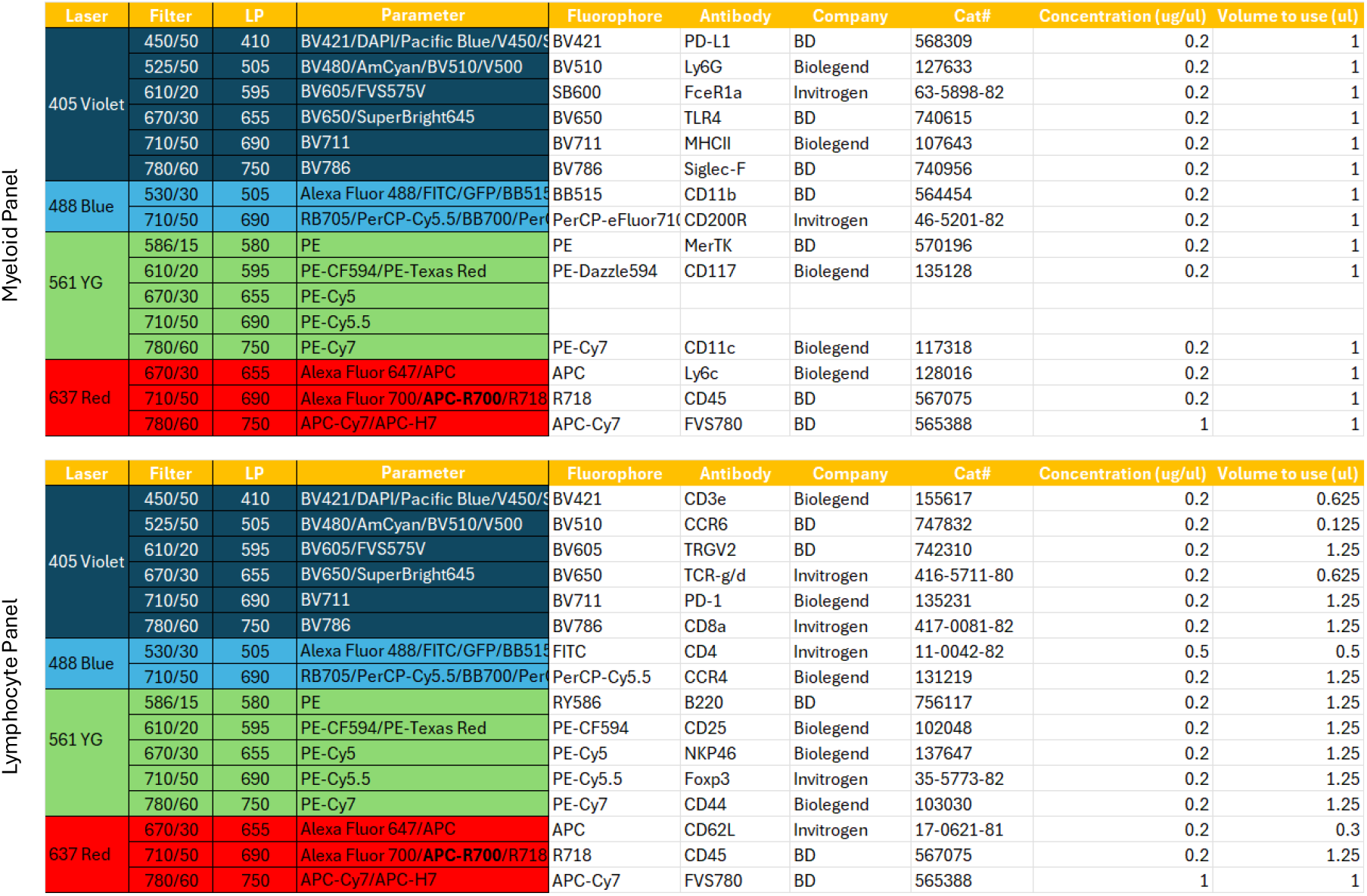
Flow cytometry panel design and information on antibodies used.

**SFig. 3.**
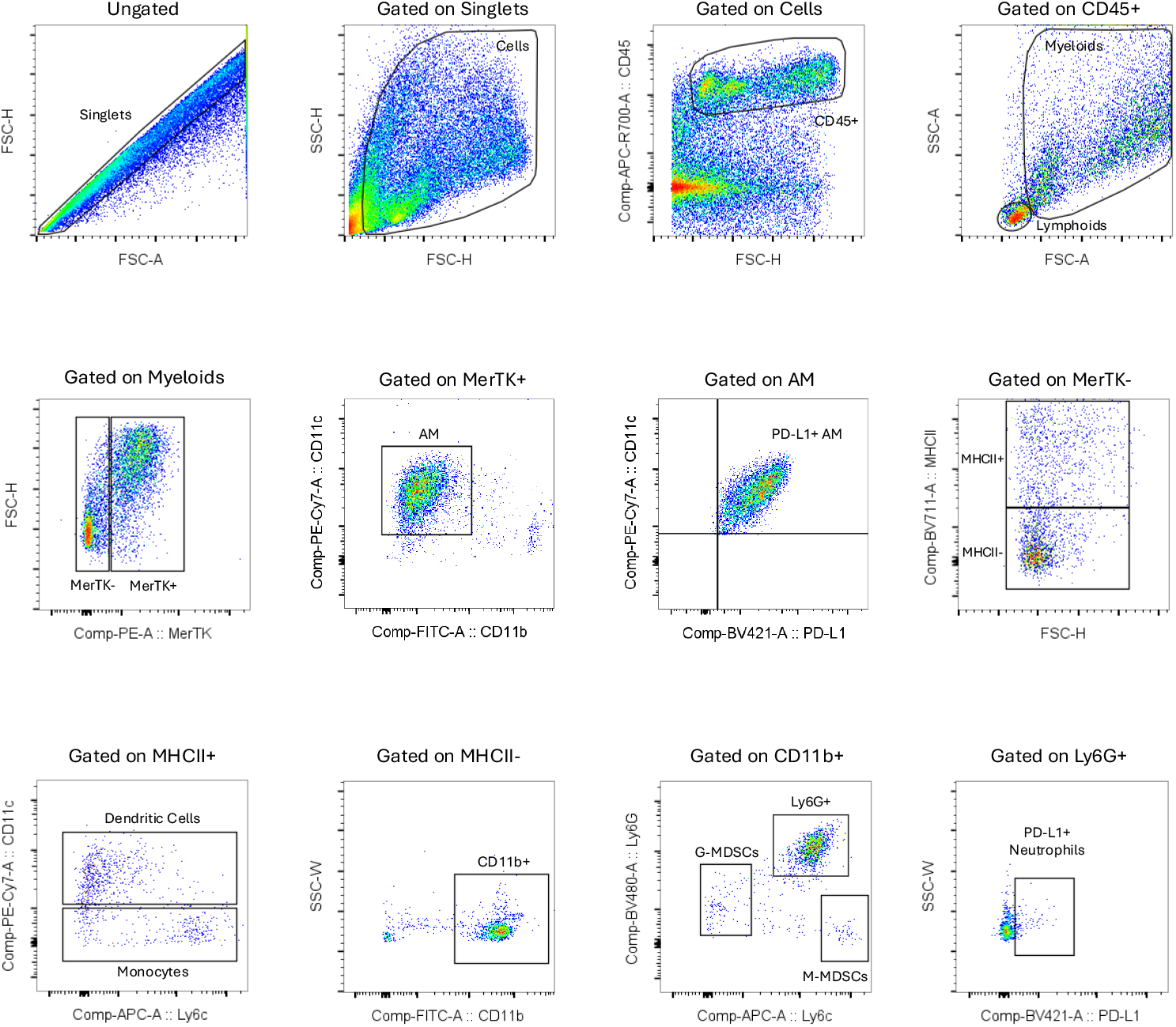
Gate strategy for myeloid cell flow cytometry analysis.

**SFig. 4.**
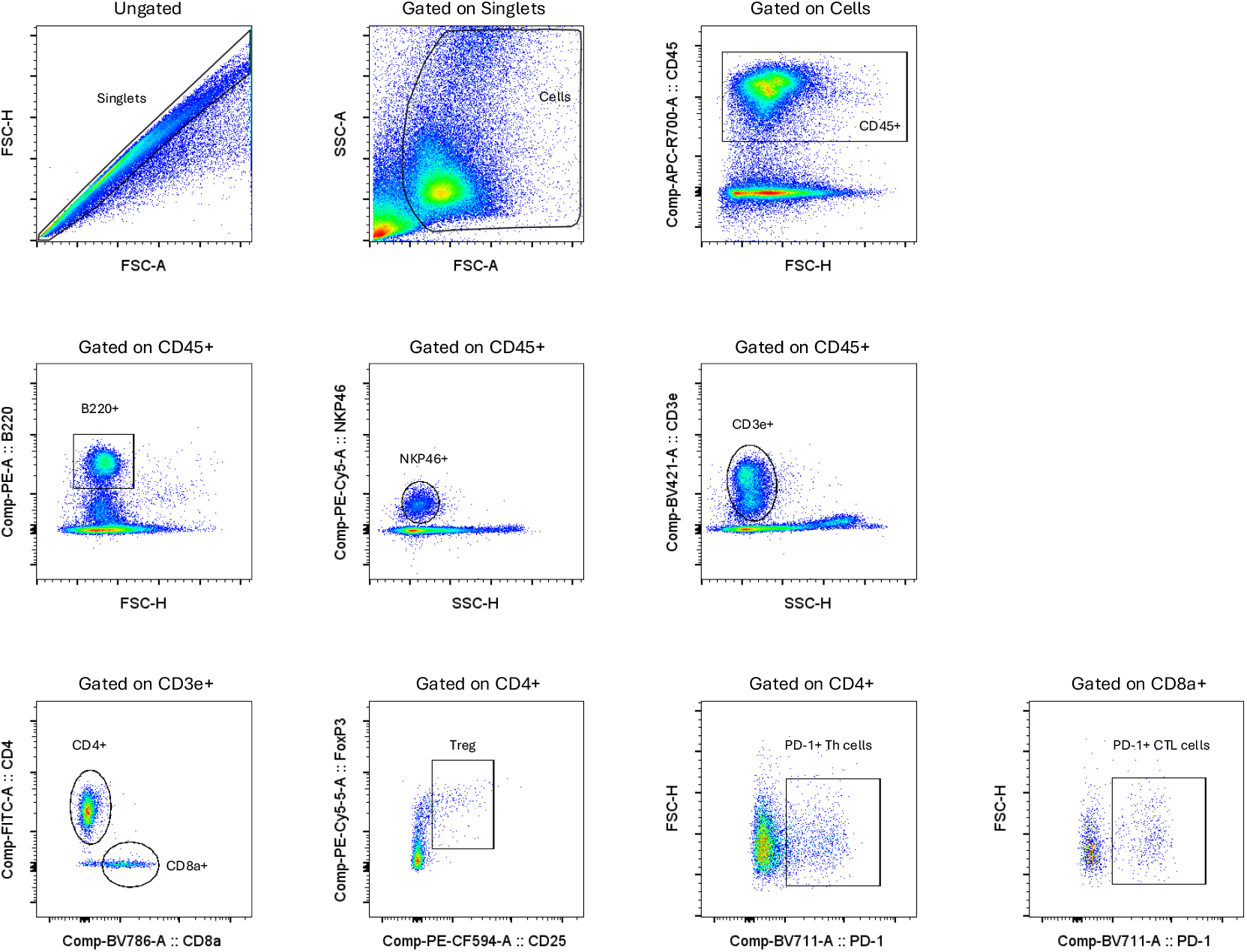
Gate strategy for lymphoid cell flow cytometry analysis.

